# Genetic introgression and transcriptomic plasticity are associated with enhanced *Leishmania infantum* pathogenicity causing human cutaneous leishmaniasis in Tunisia

**DOI:** 10.64898/2026.06.20.733521

**Authors:** Ana Maria Murta Santi, Blaise Li, Laura Piel, Juliana Pipoli Da Fonseca, Delphine Bacq-Daian, Robert Olaso, Jean François Deleuze, Thomas Cokelaer, Karim Aoun, Aida Bouratbine, Gerald F. Späth

## Abstract

The protozoan parasite *Leishmania infantum* exhibits significant genetic variability among isolates, influencing disease manifestation and treatment response. Although *L. infantum* is classically described as the causative agent of Visceral Leishmaniasis (VL) – often associated with immune deficiency, cases of Cutaneous Leishmaniasis (CL) caused by this species in immunocompetent individuals have been reported in different countries. To investigate the molecular basis of this unusual shift in tissue tropism and pathogenicity, we applied comparative genomic and transcriptomic approaches on two canine isolates (*CanL*) and two human isolates associated with Cutaneous Leishmaniasis (*CL*) in Tunisia. While the *CanL* isolates showed close genetic similarity to the *L. infantum* reference strain (JPCM5), the *CL* isolates formed a separate, highly divergent cluster based on SNP localization and frequency, differing not only from JPCM5 but also from each other. Utilizing the metagenomics sequence classification tool Kraken, we revealed a complex hybrid nature of the *CL* isolates, showing introgression from *L. donovani* and *L. tropica*, suggesting that hybridization has played a key role in generating novel phenotypic traits. Integration of RNA-seq and DNA-seq data demonstrated that only a minority of gene expression variation within and in-between the CanL or CL groups reflected gene dosage effects due to copy number variation, while the majority of expression differences were independent of gene dosage, implying post-transcriptional regulatory mechanisms contributing to parasite adaptation. In conclusion, our study identifies hybridization, genome instability, and transcriptomic adaptation as interconnected drivers of the *L. infantum* evolutionary potential. These mechanisms can collectively enhance parasite fitness gain, potentially explaining the emergence of cutaneous disease forms in a species traditionally linked to visceral infection.

**Author Summary:** This research reveals that hybridization between distinct *Leishmania* parasite species could be a key mechanism driving the evolution of new disease forms. By demonstrating that cutaneous leishmaniasis (CL) cases are caused by hybrid *L. infantum* parasites whose genomes show introgression with DNA from *L. donovani* and *L. tropica*, this study reveals a molecular mechanism potentially linked to the emergence of tegumentary disease from a species traditionally known to cause visceral infection. These findings contribute to our understanding of *Leishmania* evolution, the emergence of atypical forms of leishmaniasis linked to hybridization, and the impact of genome instability and transcriptomic adaptation as potent forces for generating phenotypic diversity and enhancing parasite fitness.

## INTRODUCTION

Leishmaniasis remains a significant global public health concern, with endemic transmission reported in over 90 countries across Asia, Africa, the Middle East, Latin America, and southern Europe. An estimated 12 million people are currently affected worldwide, while over 1 billion individuals are at risk due to exposure to the sand fly vector [1]. The disease manifests in four major clinical forms: cutaneous leishmaniasis (CL), visceral leishmaniasis (VL), mucocutaneous leishmaniasis, and post–kala-azar dermal leishmaniasis (PKDL). Global annual incidence estimates range from 600,000 to 1.2 million cases for CL and 50,000 to 100,000 cases for VL, with the highest burdens observed in countries such as Brazil, India, China, Ethiopia, Kenya, Nepal, Somalia, Sudan, Afghanistan, Bangladesh, Saudi Arabia, Syria, and Peru. The total disease burden is estimated at approximately 2.4 million disability-adjusted life years (DALYs), resulting in around 70,000 deaths annually across all forms of leishmaniasis [2].

The epidemiology of leishmaniasis is highly dynamic, depending on ecological and socioeconomic risk factors. While recurrent outbreaks are often linked to poverty, malnutrition, migration, poor sanitation or lack of medical care, disease emergence has also been associated with anthropogenic environmental effects (e.g. deforestation, climate change and population displacement). The rise for example of VL and CL cases in Europe and the Mediterranean correlates with increased international travel, migration, and higher vector densities, with upward trends observed in regions with previously low prevalence, such as North Italy or non-endemic countries, including Austria [3–5]. The encounter between EU-autochthonous and exotic, allochthonous *Leishmania* species, along with the presence of transmission-competent sand fly vectors, has been linked to the emergence of inter-species hybrids that pose a serious but often overlooked risk to public health [6].

*Leishmania* undergoes a cryptic, meiosis-like sexual cycle within the sand fly vector. Natural interspecies hybrids have been documented in regions where multiple parasite species are transmitted sympatrically [7]. Such hybrids may combine clinical and biological traits of their parental species, leading to unpredictable effects on transmission dynamics and disease outcomes. The first report of a CL outbreak involving a natural hybrid *of Leishmania infantum* (specifically, a *L. infantum/L. donovani hybrid* strain) was in the emerging focus of CL in the Çukurova region, Turkey [8]. Thousands of new CL cases have emerged in this region since 1985 and while it was originally suggested that these were due to *Leishmania tropica* infection, the causative agent has been surprisingly identified as *L. infantum*. Cases were exclusively cutaneous with no reported cases of visceral leishmaniasis (VL) in the area [9]. Whole-genome sequencing of parasite isolates from a patient and sand flies revealed that these isolates are derived from a single cross of two diverse strains with subsequent recombination within the population [8]. We recently reported a similar scenario in the Italian region of Emilia-Romagna, where an alarming increase in autochthonous visceral and tegumentary *L. infantum* infections among immunocompetent individuals has been observed [4]. Notably, this rise in human cases did not coincide with a corresponding increase in canine leishmaniasis. Instead, parasite DNA was detected in the spleens of roe deer, wild boar, and red foxes [10], suggesting a potential role of wildlife in sustaining this re-emerging focus of leishmaniasis. Microsatellite profiling and comparative genomic analyses of three strains isolated from immunocompetent visceral leishmaniasis patients revealed significant genetic introgression of the *L. infantum* genome with *L. donovani*–specific sequences [6], confirming their interspecies hybrid nature.

Similar atypical *L. infantum* strains causing CL have been observed across the Maghreb and in particular Tunisia, which are characterized by a MON-24 zymodeme and a marked polymorphism in the cysteine protease B (cpb) gene family [11,12]. However, despite their clinical relevance, comparative genomic analysis of such *L. infantum* isolates recovered from patients with CL in Tunisia are missing.

Here we fill this gap and investigate the molecular basis of this unusual shift in tissue tropism and pathogenicity applying genomic and transcriptomic approaches on two *L. infantum* CL isolates from Tunisia that were compared to two canine isolates from the same region as controls. While the CanL isolates were genetically similar to the reference strain JPCM5, the CL isolates were genetically very different from the reference showing significant introgression from *L. donovani* and *L. tropica* as judged by metagenomic analysis. Integration of RNA-seq and DNA-seq data demonstrated that gene expression variation within and between the CanL or CL groups are largely independent of gene dosage, implying post-transcriptional regulatory mechanisms contributing to parasite adaptation. Together our data suggests that hybridization is a key driver of genomic adaptation generating novel phenotypic traits that may be further fine-tuned by changes in mRNA turn-over. These mechanisms collectively enhance parasite adaptability, potentially explaining the emergence of cutaneous disease forms in a species traditionally linked to visceral infection.

## METHODS

### *Leishmania* isolates and species identification

We included in this study four *L. infantum* isolates collected from two human patients with confirmed CL cases (CL1 and CL2) and two dogs (CanL1 and CanL2) across four Tunisian regions (Figure 1A and Table 1). All isolates are included in the LR 16-IPT-06 *Leishmania* collection at Institut Pasteur of Tunis. The clinical strains were isolated on NNN medium from patients by dermal scraping, and peripheral blood of infected dogs. The four *Leishmania* isolates were identified as *L. infantum* by ITS1 PCR-RFLP and ITS1 sequencing [13].

**Figure 1:**
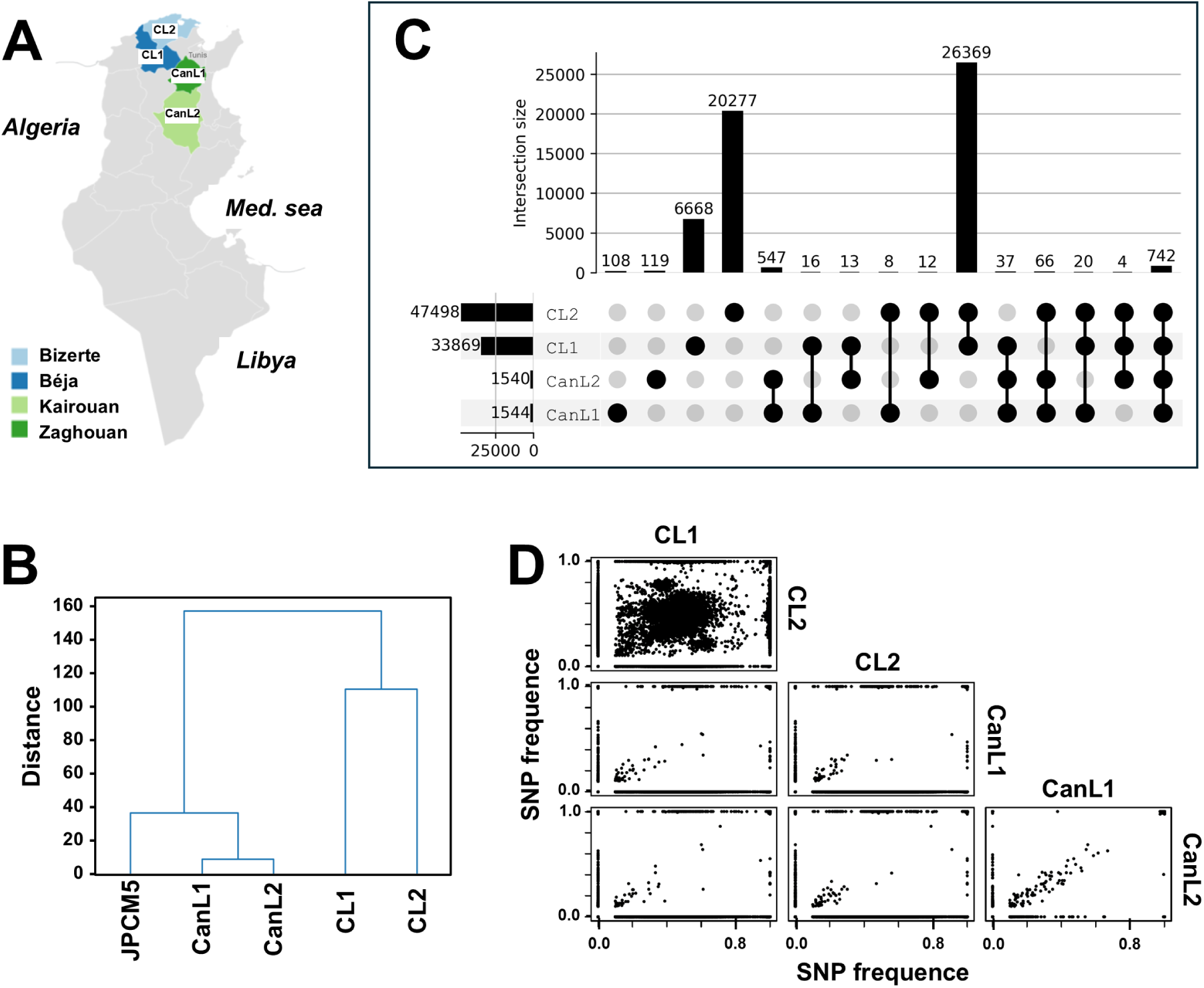
Isolation, genome sequencing and cluster analysis of four Tunisian *L. infantum* field isolates. **(A)** Geographic distribution of clinical isolates. **(B)** Cluster analysis based on distance in the SNP allele frequency space (Distance, y-axis), which quantifies the genetic dissimilarity between samples based on their allele frequencies at multiple SNP loci. **(C)** Upset plot of SNPs. The dots indicate the sample combinations. The bars in the upper histogram plot visualize the number of SNPs (indicated above the bar) common to a given combination of samples. The bars in the left histogram plot visualize the total number of SNPs (indicated to the left of the bar) for each sample. **(D)** Scatter plot showing the SNP frequence distribution between samples. Each dot corresponds to one SNP.

**Table 1:**
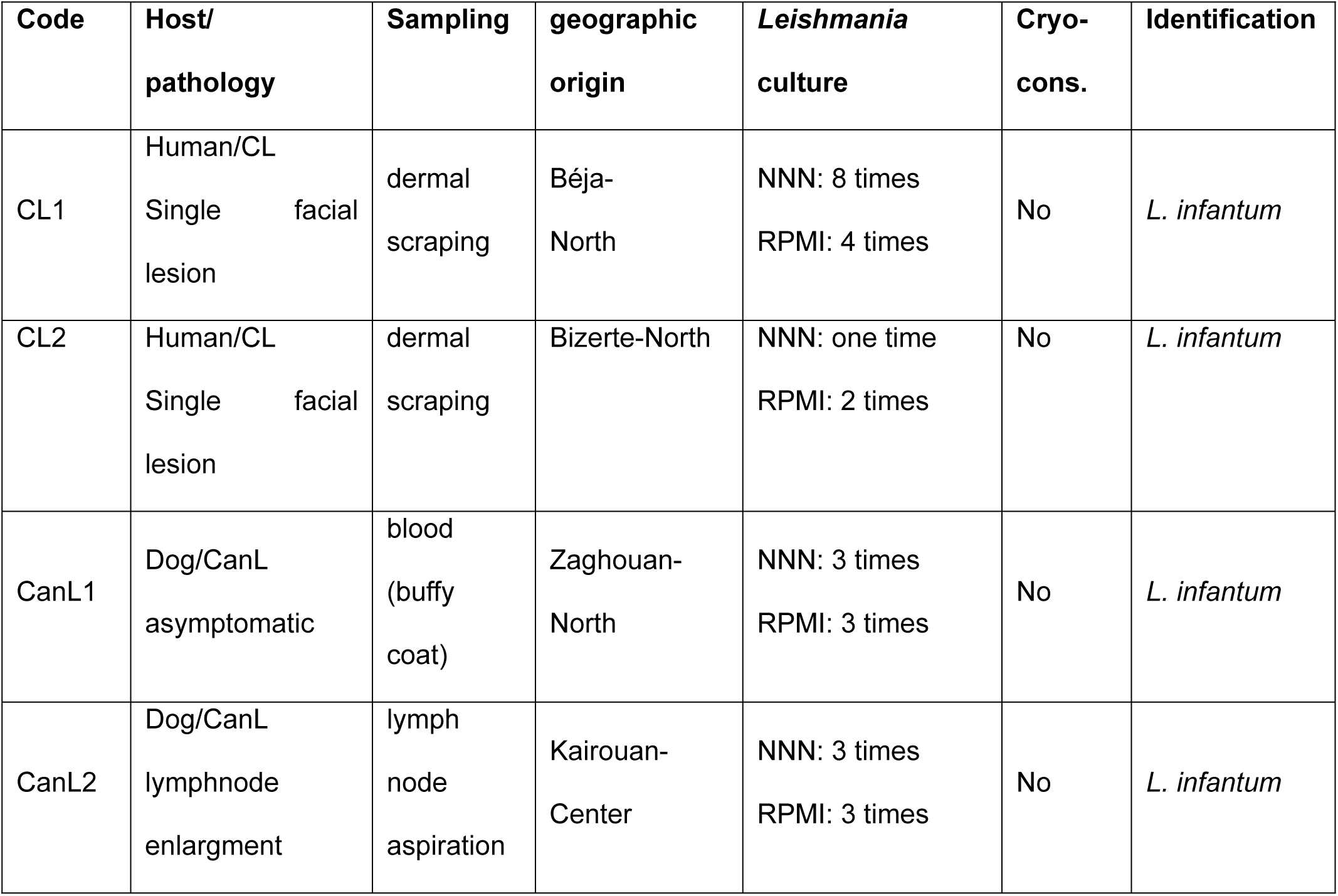
Sample description.

### DNA extraction and new generation sequencing (NGS)

Whole genome sequencing was performed by the Centre National de Recherche en Génomique Humaine (CNRGH, Institut de Biologie François Jacob, Evry, FRANCE). All concentrations of genomic DNA were measured in fluorescence (Quant-IT kits, Thermo Fischer Scientific), in duplicate, using a Molecular Device fluorescence plate reader. All samples with initial heterogenous measurements were quantified a second time and all samples with initial high concentration values were diluted to concentrations compatible with library preparation conditions and quantified again after dilution. The quality of the DNAs has been assessed on all samples by loading samples (20 ng) on a TapeStation 4200 (Agilent), in order to determine their DNA Integrity Number (DIN). After quality control, genomic DNA (1µg) was used to prepare a library for whole genome sequencing on an automated platform, using the Illumina “TruSeq DNA PCR-Free Library Preparation Kit”, according to the manufacturer’s instructions. After normalization and quality control, qualified libraries have been sequenced on a NovaSeq6000 platform from Illumina (Illumina Inc., CA, USA), as paired-end 150 bp reads. 48 samples were pooled on SP flow cell. Sequence quality parameters have been assessed throughout the sequencing run and standard bioinformatics analysis of sequencing data was based on the Illumina pipeline to generate a FASTQ file for each sample.

### DNAseq analysis

WGS reads were mapped against the *L. infantum* reference genome JPCM5 [14] and sequencing data processed using GIP version 1.1.0 [15] (see Table S1 for sequencing statistics). The outputs of the GIP pipeline were used for downstream analyses, ignoring the reads mapping to mitochondrial DNA (i.e. maxi and minicircles). Mean coverage in 300 bp bins as generated by the GIP pipeline were used to compute somy scores per chromosome by first normalizing bin scores for a sample by their median across the entire genome (to obtain comparable values between samples), multiplying by two (to scale somy values to the default diploid state assumed for most of the chromosome). These somy scores were represented on ‘per chromosome and per sample’ boxplots. The median somy score across bins belonging to a given chromosome were visualized on a sample versus chromosome heatmap.

Normalized mean coverages per gene are reported by GIP, as the mean coverage of the gene divided by the median coverage of the chromosome containing the gene. These values were used to compute a heatmap, as follows. Gene coverage tables generated by GIP were loaded in a Jupyter notebook [16] using the Pandas (version 1.4.2) [17,18] Python library. Genes that exhibited less than 1 unit difference in normalized mean coverage between all pairs of samples were discarded, as well as those where the average MAPQ score was below 10 for at least one of the samples. Scaled normalized mean coverages were computed (subtracting the mean across samples and dividing by the standard deviation, using the preprocessing.scale function from the scikit-learn (version 1.0.2) Python library [19], with options with_means and with_std set to True) and used as input for the clustermap function of the seaborn (0.11.2) [20] Python library. Clustering of the samples was deactivated (option row_cluster set to False).

SNP frequencies were retrieved from the filtered output of GIP for each sample. For a given set of samples, the union of all SNPs across samples was considered, assuming an alt-allele frequency of 0 when a SNP was missing from a given sample. Distributions of alt-allele frequencies were represented as upset plots generated using the upsetplot 0.6.0 Python library [21], considering SNPs at alt-allele frequency 0 as missing from a given sample. SNPs were sorted according to genomic coordinates and assigned a corresponding index against which alt allele frequencies were plotted. Euclidean distances between samples in the SNP allele frequencies space were computed and used to build a dendrogram using the scipy.cluster.hierarchy module from the Scipy 1.8.<u>0</u> Python library [22]. The clustering was obtained using the UPGMA method. Besides the more specialized Python libraries indicated above, post-GIP analyses were carried out using the Pandas 1.4.2, Matplotlib 3.5.1 and Seaborn 0.11.2 Python libraries. Taxonomic assignation of raw Illumina reads for some of the samples was performed using kraken 2.1.3 [23,24] against the “protozoa” database to get insights into the possible hybridization events at the origin of putative hybrid samples. The summarized taxonomic assignations were extracted from the reports generated by kraken and represented as Sankey plots, using the plotly (6.1.0) Python library [25].

### RNA extraction and sequencing

Total RNA was extracted from promastigotes in exponential culture phase. Promastigotes were centrifuged at 2,000 x *g* for 10 min at 4°C and re-suspended in the lysis buffer supplied with the NucleoSpin RNA Plus (macherey-nagel). The samples were stored at −80°C and RNA extractions were performed according to the manufacturer’s instructions, including a DNase treatment. RNA integrity was controlled using the Agilent Bioanalyzer. DNase-treated RNA extracts were used for library preparation using the Illumina Stranded Total RNA prep and custom ribodepletion probes designed using the *L. infantum* reference genome JPCM5 (Illumina, San Diego, California) according to the manufacturer’s instructions. An initial ribodepletion step (included in the Illumina protocol) was performed on total RNA to remove ribosomal RNA. Fragmentation was performed on the enriched fraction by divalent ions at high temperature. The fragmented RNA samples were randomly primed for reverse transcription followed by second-strand synthesis to create double-stranded cDNA fragments. No end repair step was necessary. An adenine was added to the 3’-end and specific Illumina adapters were ligated. Ligation products were submitted to PCR amplification. The obtained oriented libraries were controlled by Fragment Analyzer NGS high sensitivity (Agilent, DNF-474-1000) and quantified by spectrofluorimetry (Quant-iT High-Sensitivity DNA Assay Kit, #Q33120, Invitrogen). Sequencing was performed on the Illumina Novaseq platform at the Biomics platform (Institut Pasteur, Paris, France) to generate paired end, 150-bp reads bearing strand specificity.

### RNA-seq analysis

The bioinformatics analysis was performed using the Sequana RNA-seq pipeline (version 0.18, https://github.com/sequana/rnaseq) from the Sequana project [26]. Reads were cleaned of adapter sequences and low-quality sequences using fastp software (version 0.23) [26]. Bowtie2 (version 2.5.4) [27], with default parameters, was used for alignment on the reference genome (*Leishmania infantum* from TritrypDB v61). Genes were counted using featureCounts (version 2.0.1) [28] from Subreads package (parameters: -t gene -g gene_id -s 2). Count data were analyzed using R version 4.1.2 [29] and the Bioconductor package DESeq2 (version 1.34.0) [30]. The normalization and dispersion estimation were performed with DESeq2 using the default parameters and statistical tests for differential expression were performed applying the independent filtering algorithm. For each pairwise comparison, raw p-values were adjusted for multiple testing according to the Benjamini and Hochberg (BH) procedure [31] and genes with an adjusted p-value lower than 0.05 were considered differentially expressed.

### Gene Ontology Enrichment Analysis

To identify biological processes significantly overrepresented within each of our gene sets, Gene Ontology (GO) enrichment analyses were performed. The analyses were conducted using the GO Enrichment tool available on TriTrypDB (https://tritrypdb.org/tritrypdb/app/), release 68 (7 May 2024) [32,33]. The following parameters were used for the analysis: *L. infantum* JPCM5 as the reference organism, the Biological Process ontology, Computed and Curated evidence, without limitation to GO Slim terms, and a p-value cutoff of 0.05. The background population for the statistical test was the entire set of genes for *L. infantum* JPCM5 available in the database. The statistical significance of GO term enrichment was determined using a Fisher’s exact test, with a p-value cutoff of 0.05. To correct multiple hypothesis testing, both the Benjamini and Bonferroni correction methods were applied. Enriched GO terms were identified as those present in the gene subset at a frequency significantly higher than in the background set of all genes for the organism.

## RESULTS

### *L. infantum* strains causing human CL are genetically distinct from canine isolates

We performed comparative genome and transcriptome analyses of four *L. infantum* field isolates collected from distinct Tunisian governorates (Figure 1A and Table 1). Two isolates (CL1 and CL2) were obtained via dermal scraping from facial lesions of immunocompetent patients with acute, cutaneous leishmaniasis. The other two isolates (CanL1 and CanL2) were collected from infected dogs - one asymptomatic and the other showing lymph node enlargement. To minimize culture-induced genetic changes associated with in vitro adaptation [34], DNA was extracted from short-term cultured parasites (Table 1), subjected to Illumina short-read sequencing, and reads were aligned to the *L. infantum* reference genome JPCM5 [14] (see Table S1 for sequencing metrics).

We applied our genome instability pipeline (GIP) and associated *giptools*, which allows the batch analysis of multiple genomes, including mapping of short reads, quantification of chromosomes, genes and genomic bins [15]. Using GIP we first assessed evolutionary divergence among the isolates through cluster analysis of allele frequency differences at multiple SNP loci. The canine isolates clustered closely with JPCM5, indicating minimal divergence from the reference genome. In contrast, CL1 and CL2 formed a distinct, deeply branched cluster, reflecting significant divergence not only from JPCM5 but also between the two human-derived isolates themselves (Figure 1B).

While only around 1,500 SNPs were detected in CanL1 and CanL2 (over 90% or 1,392 shared SNPs), CL1 and CL2 showed respectively 33,869 and 47,498 SNPs compared to the *L. infantum* reference genome (Figure 1C). Of these, 27,135 SNPs were shared by both CL isolates, while 6,668 (19%) and 20,277 (42%) SNPs were unique to CL1 and CL2, respectively. This contrast between the genetic similarity of CanL samples and the divergence of CL isolates is further supported by pairwise correlation plots of SNP frequencies (Figure 1D): SNPs in CanL samples fall largely along a diagonal, indicating highly similar allele frequencies, whereas SNPs shared between CL1 and CL2 are dispersed across the plot, highlighting substantial allele frequency differences that may suggest allelic selection.

In conclusion, based on SNP number and frequency, the two human *L. infantum*isolates appear to share an ancient common ancestry but have since undergone divergent evolution. Despite the limited sample size, the distinct genetic and clinical profiles of the four isolates offer a valuable opportunity to investigate in the following the genomic and transcriptomic signatures linked to the pathogenicity of CL1 and CL2.

### The human *L. infantum* isolates are characterized by genetic introgression

Genetic introgression - the transfer of genetic material from one species into the gene pool of another through hybridization [35] - was evident in the CL isolates when compared to the CanL isolates. Analysis of SNP frequency and localization using the *giptools* SNV module [15] revealed strong signals of heterozygosity in CL1 and CL2, with allele frequencies largely centered around 50% (Figure 2A and B, Figure S1B). This pattern is typically indicative of hybridization, with each allele inherited from a different parental species.

**Figure 2:**
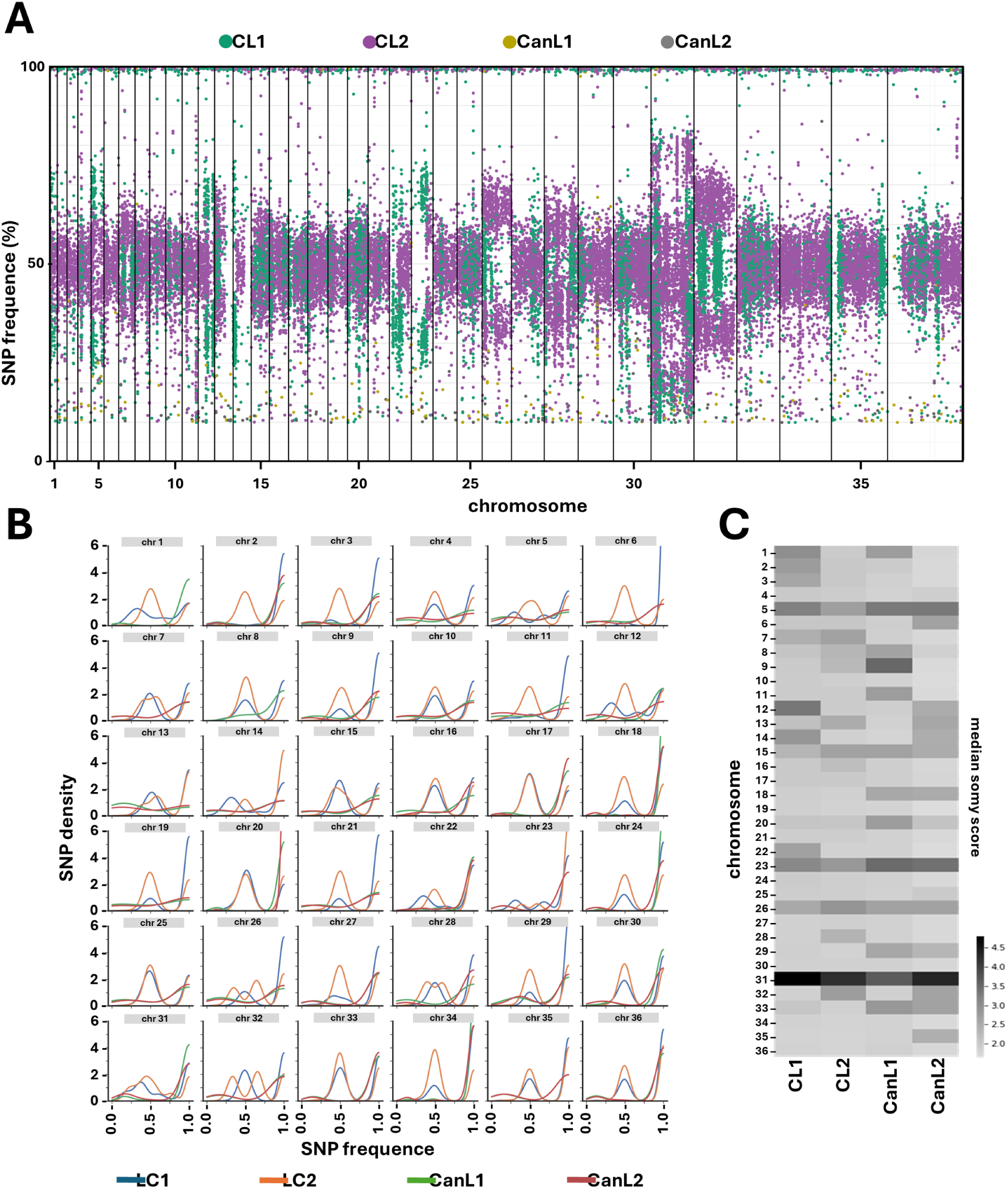
Genome-wide SNP analyses. **(A)** SNP genomic localization. The x-axis indicates the genomic position while the y-axis indicates the SNP frequency. Each dot corresponds to one SNP. The different chromosomes are indicated, and their boundaries are visualized as vertical lines. The individual SNPs are colored according to the samples as indicated in the legend. **(B)** SNP density distribution. Estimated probability of SNP densities (y-axis) are plotted as a function of SNP frequence (x-axis). Different chromosomes are reported in different panels and lines are colored according to sample as indicated in the legend. **(C)** Heatmap of the median chromosome somy scores of the indicated samples.

The heterozygous SNPs were not uniformly distributed across the genome but localized in chromosomal sub-regions. Such genetic disequilibrium likely results from repeated backcrossing or gene conversion events, both of which can disrupt the original balance between parental alleles leading to localized loss of heterozygosity. The SNP accumulation in defined chromosomal regions only partially overlapped between CL1 and CL2, further supporting their divergent evolution from a putative, shared hybrid ancestor. Alternatively, this SNP pattern could also be explained by independent hybridization events followed by convergent selection towards a similar genome structure.

The heterozygous SNPs are defining two distinct haplotypes, which can be phased for certain aneuploid chromosomes by haplotype specific frequency shifts from 50:50 to 33:66 (e.g. chr5, 12, 22, or 23 for CL1; and chr13, 26 or 32 for CL2) (Figure 2A and B). Estimating the somy score (see methods) as a proxy for ploidy, we observed converging amplification across all isolates for chromosomes 5, 15, 23, and 26 (Figure 2C and Figure S1A), some of which were indeed linked to haplotype-specific frequency shifts. Amplification of these chromosomes has been previously associated with in vitro fitness gain in Indian and Sudanese *L. donovani* strains [36–39], suggesting that these karyotypic changes may result primarily from culture adaptation - despite the limited number of passages - rather than reflect epidemiologically significant variation.

### The human *L. infantum* isolates represent a novel class of hybrids characterized by a complex, mosaic genome architecture

Cluster analyses based on SNP allele frequency placed CL1 and CL2 in a branch that contained the two previously published, atypical Tunisian human isolates ZK28 and LIPA83 [40], various canine samples from Tunisia and the JPCM5 reference genome (Figure 3A). CL1 and CL2 are thus clearly distinct from previously published *L. infantum* hybrids linked to visceral infection in Italy (leish7, leishMO23, leishMO38), Cyprus (CH isolates) and Turkey (CUK isolates) (Figure 3A) [6,8,41].

**Figure 3:**
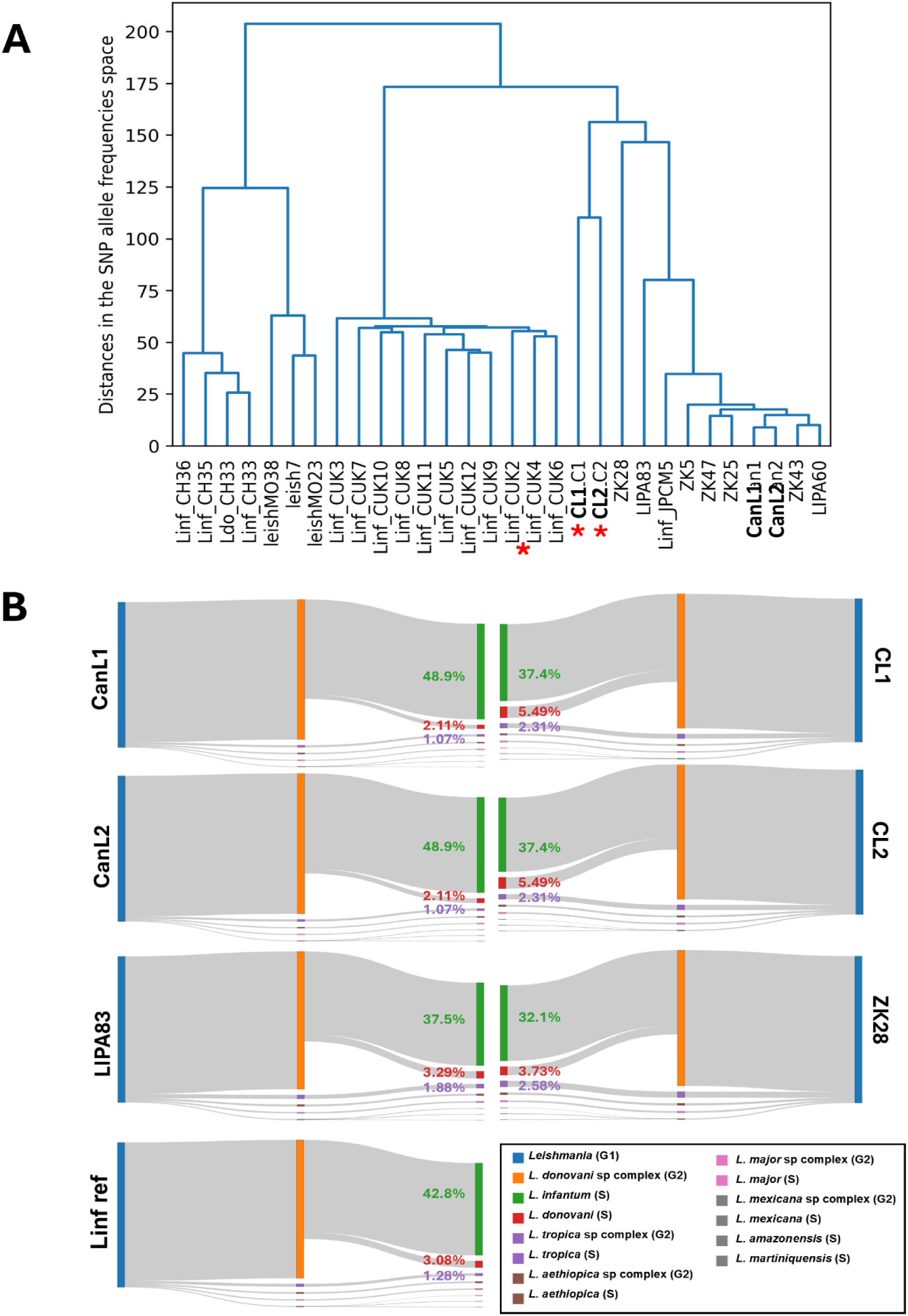
Analysis of genome structures. **(A)** Cluster analysis based on pairwise Euclidean distances between alt allele frequencies across SNPs. The tree is constructed from those distances using the UPGMA method. Some samples that were not sequenced in the present study have their names prefixed by a tag indicating the species: Ldo, *L. donovani*; Linf, *L. infantum*. The samples that were sequenced in the present study are indicated by a red asterisk. **(B)** Sankey plots visualizing the results of the Kraken analysis. The shown percentages correspond to the reads unambiguously and uniquely assigned for each sample to a given taxon as indicated by the color code (see legend).

To further investigate the hybrid nature of CL1 and CL2, we utilized the Kraken sequence classification tool, which is optimized for metagenomic analyses and assigns taxonomic labels to DNA sequences through exact k-mer alignment [23,24]. Although the majority of reads across all samples were classified within the *L. donovani* species complex (Figure 3B), the proportion of uniquely assigned reads at this taxonomic level was over 10% lower in human-derived samples compared to canine samples, consistent with the presence of genetic introgression. Indeed, in the human samples, over 5 % and 2.4% of reads were uniquely assigned to respectively *L. donovani* and *L. tropica,* which was more than double those observed in canine samples, with <2.5% for *L. donovani* and <1.1% for *L. tropica*). The atypical *L. infantum* strains LIPA83 and ZK28 showed a pattern similar to the *L. infantum* reference strain, which fall between the CL and CanL values for the *L. donovani* and *L. tropica* k-mer assignment.

Together, these data reveal a complex structure of the *L. infantum* CL strains containing genetic introgression from both *L. donovani* and *L. tropica* caused by previous intra-species hybridization events, followed by possible gene conversion events during self-hybridization (i.e. selfing) [42].

### Gene dosage-independent expression changes are the major driver of parasite evolution in the field

Conceivably, the genetic introgression we observed in the CL isolates is the result of previous hybridization events, which initially should generate aneuploid offspring as observed previously [43,44]. A recent report has revealed extensive genomic and transcriptomic remodeling in inter-species hybrids that can compensate for the genomic shock caused by the fusion of different parental genomes [45]. We next assessed to what extent the CL hybrid isolates mitigated this genomic shock undergoing genomic and transcriptomic adaptation.

In contrast to karyotypic changes (see Figure 2C), which are highly dynamic and typically associated with early culture adaptation, gene copy number variations (CNVs) tend to be more stable and can therefore provide insights into field-relevant genomic adaptations, particularly when analyzing parasites during the early stages of in vitro cultivation [38]. The heatmap in Figure 4A shows the normalized mean gene coverage across all samples, generated with the *geInteraction* module of *giptools [15]*. The canine and human isolates form distinct clusters, each characterized by unique gene copy number variation (CNV) profiles (Figure 4A). Notably, both the canine and human samples exhibit increased read depth across numerous genes previously associated with parasite virulence, including members of the surface antigen protein 2 family (LINF_120013500, cluster 1; LINF_120014700, cluster 2), or various amastins (LINF_080012000, cluster 1; LINF_340034900 / LINF_340010400 / LINF_340010600, cluster 3; LINF_340024000, cluster 4) (Table S2). While clusters 1 and 2 indicate that some gene CNVs are under convergent selection in either the canine or the human samples, clusters 3 and 4 reveals gene CNVs that are distinct within each sample set, suggesting strain-specific, divergent evolution.

**Figure 4:**
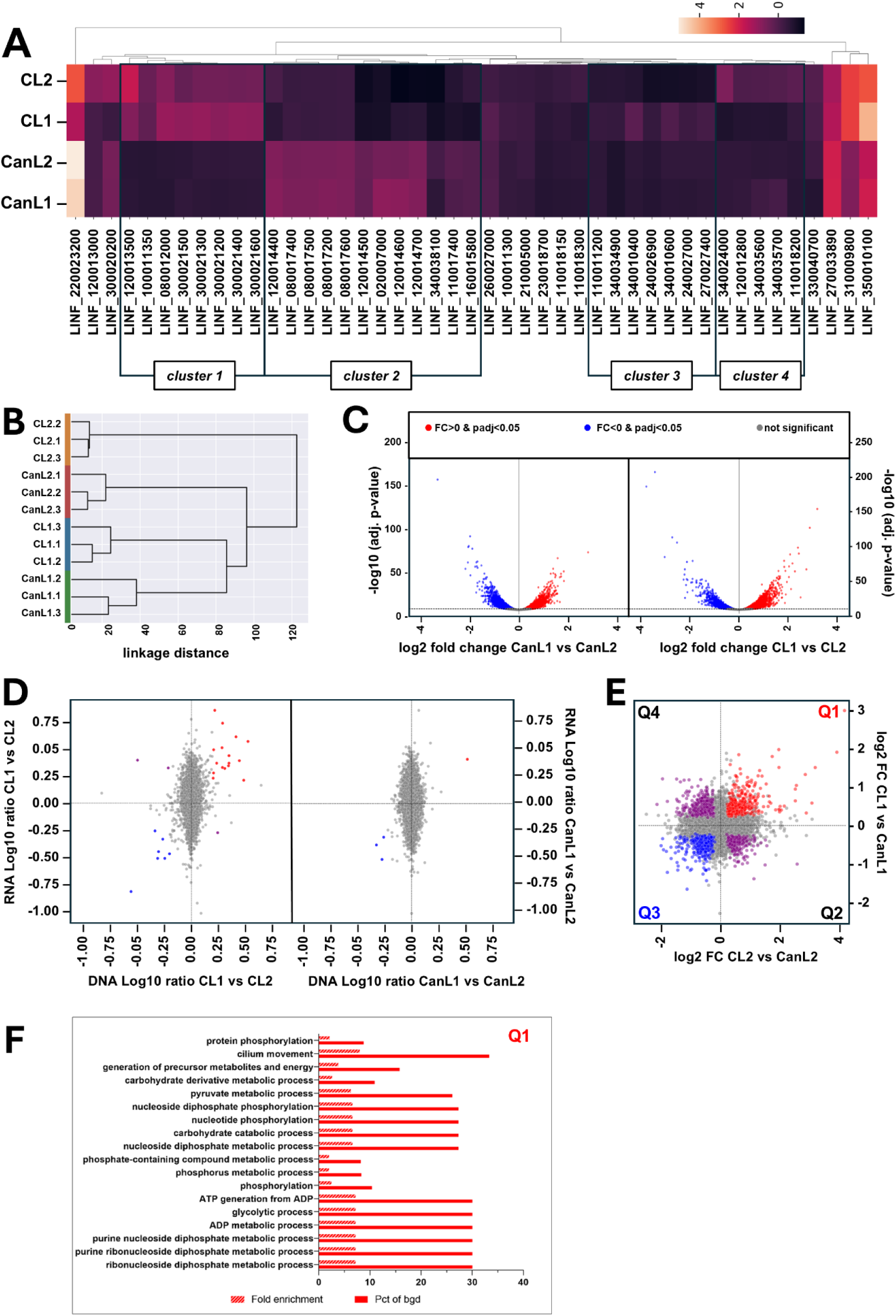
Multi-omics comparison of genomic and transcriptomic alterations in human versus canine isolates. **(A)** Heatmap of gene copy number variations between the four isolates identified with the geInteraction module of the GIP package [15]. Only reads with MAPQ>10 were considered. **(B)** Hierarchical clustering of the RNAseq sample set. The expression data were log-transformed before computing the Euclidean distance between samples and building the dendrogram using the Ward method. The numerical values shown in the dendrogram (horizontal axis) represent the linkage distances or dissimilarities at which clusters are merged during the clustering process. **(C)** Volcano plots of RNAseq data comparing the two canine (left) and two human samples (right). Grey dots correspond to transcripts whose changes in abundance did not meet the significance criteria (i.e. adjusted p-value > 0.05). **(D)** Double ratio plots showing RNA and DNA read depth ratios between the two human (right) and two canine samples (left). Red, increased read depth ratios for both DNA and RNA; blue, decreased read depth ratios for both DNA and RNA; purple, counter-correlated read depth ratios; grey, DNA read depth ratios < 1,5. (E) Double ratio plots showing DNA-normalized RNA read depth ratios. Red, transcripts with increased abundance in both comparisons CL1/CanL1 and CL2/CanL2; blue, transcripts with decreased abundance in both comparisons CL1/CanL1 and CL2/CanL2; purple, transcripts showing counter-correlated changes in abundance; grey, non-significant changes (adjusted p-value < 0.05). (F) Bar plots showing enriched GO terms for biological processes identified in quadrants Q1 (left) and Q3 (right).

In parallel to genome sequencing, we performed RNA-seq analysis of the canine and human samples, for each using three independent cultures derived from individual isolates. Principle component analysis (Figure S2A) revealed only little variation between the replicates, but important variation between isolates, indicating that transcriptomic differences largely override genetic divergence. Surprisingly, the human and canine isolates did not cluster with respect to host origin (Figure 4B), suggesting important transcriptomic divergence within and in-between strains. Indeed, pairwise comparisons CanL1_vs_CanL2 and CL1_vs_CL2 (Figure 4C) demonstrated that approximately 50% of all genes showed significant expression changes (Table S3).

Integrating the DNA- and RNA-seq datasets confirmed gene dosage-dependent expression changes as judged by the (weak) correlation between chromosome/gene copy number variations and transcript output (Figure S2B). However, double ratio plots comparing DNA and RNA read depth variations within the canine or human samples (Figure 4D) reveals that most expression changes occur in a gene dosage-independent fashion, with the dynamic range of RNA abundancies varying between +5 (Log10=0.75) and -10fold (Log10=-1) for genes that show constant DNA read depth across samples. Direct comparison of DNA-seq-normalized RNA-seq results between canine and human samples (CL1 vs CanL1 to CL2 vs Lcan2, Figure 4E) informs on post-transcriptional changes that converge within the two human isolates (quadrants Q1 for increased and Q3 for decreased expression) when compared to the two canine samples.

Gene Ontology enrichment analysis was conducted on the set of transcripts commonly upregulated in both CL samples relative to the CanL group (Q1, 364 gene IDs). After applying the Benjamini-Hochberg procedure for multiple testing correction (FDR < 0,05) (Table S4), GO terms for 19 biological processes were significantly enriched, centered around three primary themes: energy metabolism and ATP generation (e.g., glycolytic process, ATP generation from ADP), phosphorus and phosphate metabolic processes (e.g., phosphorylation), and cellular motility (cilium movement) (Figure 4F). Several terms show exceptionally high fold enrichment (≥ 7), including the GO terms ‘nucleoside diphosphate/ATP metabolic processes’ and ‘glycolysis’, indicating these pathways are highly specific to the gene set. On the other hand, no GO terms showed significant enrichment among the transcripts commonly downregulated in both CL samples compared to CanL (Q3, 301 gene IDs) after correcting for multiple testing (Table S4).

In conclusion, both genome instability and transcriptomic reprogramming participate in mitigating potential toxic effects associated with *Leishmania* hybridization and are key drivers of parasite adaptation and fitness gain in the field.

## DISCUSSION

A critical yet often overlooked factor contributing to the emergence and adaptability of *Leishmania* is the generation of inter-species hybrids through a cryptic, sexual cycle within the sand fly vector [43,44,46–51]. In a recent report, we linked emerging VL in northern Italy to a *L. infantum/L. donovani* hybrid population that preferentially infects humans rather than dogs [6]. Here we identify a new class of putative *L. infantum* hybrids in Tunisia, showing genetic introgression from *L. donovani* and *L. tropica*, and associated with cutaneous disease. The emergence of these strains, combined with their unusual epidemiology, and their increased pathogenic potential, confirm hybridization as a key risk factor for *Leishmania* infection in the Mediterranean basin and Europe.

Inter-species hybridization is driven by the co-occurrence of different parasite species within the same sand fly vector [7,50]. Such sympatric transmission cycles can be fuelled by population displacement and are frequently associated with the manifestation of atypical clinical phenotypes. For example, extensive hybridization involving *L. donovani* with *L. major* and *L. tropica* causes cutaneous leishmaniasis in Sri Lanka [47,48]. Like the situation observed in Tunisia, a focus of CL outbreaks was caused in the Çukurova province of southeast Turkey by hybrids between *L. infantum* and members of the *L. donovani* species complex (CUK isolates) [8,9]. Similar scenarios have also been reported in the Americas, where *L. peruviana/L. Braziliensis* hybrids cause severe cutaneous and mucocutaneous pathologies [52]. Alterations in *Leishmania* disease patterns are not only associated with inter-species hybridization but have also been observed in an intra-species *L. donovani* hybrid responsible for CL cases in the Himachal Pradesh region of India [47]. The increased detection of hybrids in recent studies may be explained by two phenomena. On one hand, hybridization may be a rare process, but hybrids could be disproportionately detected due to their consistent fitness advantage and distinctive clinical presentation. On the other hand, hybridization may be a relatively frequent event, with independent hybrids often converging toward a similar clinical phenotype. Such convergent evolution is supported by our findings: both the Tunisian CL and Turkish CUK isolates appear to have arisen from independent hybridization events as judged by their large genetic distance (see Figure 3A). Nevertheless, both converge on a similar clinical phenotype causing disease in immunocompetent individuals, exhibiting a possible anthroponotic transmission pattern, and showing a shift from visceral to cutaneous tropism [9,53].

These examples raise an important question: how does hybridization promote evolvability in ways that can facilitate ecological expansion, broaden host range, alter tissue tropism, and drive disease emergence? Our study implicates both genomic and phenotypic mechanisms in this process. First, hybridization increases heterozygosity enabling the generation of novel, advantageous allele combinations. The presence of heterozygous SNPs is considered diagnostic for hybrid lineages. Indeed, only the CL but not the CanL isolates showed high density of SNPs with a frequence of 50%. These SNPs were localized in defined sub-chromosomal regions, indicating reduced linkage disequilibrium, likely due to gene conversion events or genetic recombination occurring during mitosis or selfing [50]. In previous studies, we have identified similar patches as hot spots of recombination during sand fly infection, allowing for haplotype shuffling and the selection of beneficial alleles [54].

Second, hybridization can cause immediate changes in gene dosage that can modulate gene expression. We and others have shown that chromosome and gene amplification correlates with increased transcript output [36,37,55–57]. Comparison of genomic and transcriptomic read depth allowed us in the current study to confirm this observation within each experimental group (Figure 4D and S2B). Significant differences in gene CNVs were observed between the CL and CanL samples, affecting known virulence factors such as surface antigen protein 2 (Figure 4A and Table S2). Conceivably, gene dosage-dependent, differential expression of these factors may participate in driving the atypical clinical manifestation observed in the CL hybrids.

Finally, aside these adaptive genomic changes, hybridization may also drive phenotypic adaptation at post-transcriptional, translational or post-translational levels. Applying our *Leishmania* experimental evolution system, we have previously observed gene dosage-independent variations in mRNA abundance associated with *Leishmania* fitness gain *in vitro*, causing for example reduced expression of flagellar components and increased expression of factors associated with accelerated proliferation in culture, such as ribosomal components or snoRNAs [38]. In addition, we have observed that not all genetic amplifications correlate with transcript levels, revealing an important role of mRNA turn-over in adaptation and compensation for toxic gene dosage effects [39]. Analysing the gene dosage-independent transcript profiles between canine and human isolates revealed important differences in mRNA expression demonstrating the relevance of phenotypic (non-genomic) adaptation in *Leishmania* evolution in clinical settings. These gene dosage-independent, transcriptomic changes converged in the human samples on GO terms linked to signal transduction, suggesting possible roles of post-translation regulation in parasite adaptation and fitness gain. These adaptive transcriptomic responses may further contribute to the differences in pathogenicity observed between human- and canine-derived *L. infantum* strains.

In conclusion, intra- and inter-species hybridization acts as a reservoir of genetic innovation that can enhance parasite fitness in natural populations. Hybrids frequently exhibit traits that surpass those of their parental lines, as recombination and gene conversion events can reshuffle virulence factors across species, often producing unpredictable effects on pathology, epidemiology, and disease outcomes. While the generation of such genetic diversity may occur randomly, the retention of parental alleles is likely non-random and may be the result of natural selection. Consequently, future studies of hybrid field isolates should integrate long-read genome sequencing with transcriptomic analyses to (i) fully resolve the mosaic architecture of hybrid genomes and trace their evolutionary origins, (ii) evaluate the contributions of phenotypic adaptation and allelic expression in the atypical epidemiology of hybrid infection, and (iii) identify genes under natural selection in these hybrids that could reveal novel pathogenicity factors with diagnostic or prognostic relevance.

## ACKNOWLEDGEMENTS

We thank all the CEA-CNRGH staff who contributed to sample preparation and sequencing for their excellent technical assistance. We thank the Institut Pasteur Biomics Platform, C2RT and in particular Georges Haustant for technical assistance.

## AUTHOR CONTRIBUTIONS

AMMS: Conceptualization, Formal analysis, Investigation, Methodology, Validation, Visualization, Writing – original draft, Writing – review & editing

BL: Data curation, Formal analysis, Methodology, Software, Visualization, Writing – review & editing

LP: Formal analysis, Visualization, Writing – review & editing

JPL: Formal analysis, Investigation, Methodology, Visualization, Writing – review & editing

DB: Conceptualization, Funding acquisition, Methodology, Project administration, Resources

RO: Conceptualization, Funding acquisition, Methodology, Project administration, Resources

JFD: Conceptualization, Funding acquisition, Methodology, Project administration, Resources

TC: Data curation, Formal analysis, Methodology, Software, Visualization, Writing – review & editing

KA: Conceptualization, Funding acquisition, Project administration, Resources, Supervision, Writing – original draft, Writing – review & editing

AB: Conceptualization, Funding acquisition, Project administration, Resources, Supervision, Writing – original draft, Writing – review & editing

GFS: Conceptualization, Data curation, Formal analysis, Funding acquisition, Methodology, Project administration, Resources, Supervision, Visualization, Writing – original draft, Writing – review & editing

## REFERENCES

1. WHO. Leishmaniasis [Internet]. 2023 [cited 2025 Nov 19]. Available from: https://www.who.int/news-room/fact-sheets/detail/leishmaniasis

2. Mann S, Frasca K, Scherrer S, Henao-Martínez AF, Newman S, Ramanan P, et al. A Review of Leishmaniasis: Current Knowledge and Future Directions. Curr Trop Med Rep. 2021 Mar 17;8(2):121–32. doi:10.1007/s40475-021-00232-7

3. Riebenbauer K, Czerny S, Egg M, Urban N, Kinaciyan T, Hampel A, et al. The changing epidemiology of human leishmaniasis in the non-endemic country of Austria between 2000 to 2021, including a congenital case. Reiner RC, editor. PLoS Negl Trop Dis. 2024 Jan 10;18(1):e0011875. doi:10.1371/journal.pntd.0011875

4. Todeschini R, Musti MA, Pandolfi P, Troncatti M, Baldini M, Resi D, et al. Re-emergence of human leishmaniasis in northern Italy, 2004 to 2022: a retrospective analysis. Eurosurveillance. 2024 Jan 25;29(4). doi:10.2807/1560-7917.ES.2024.29.4.2300190

5. Gaspari V, Gritti T, Ortalli M, Santi A, Galletti G, Rossi A, et al. Tegumentary Leishmaniasis in Northeastern Italy from 2017 to 2020: A Neglected Public Health Issue. Int J Environ Res Public Health. 2022 Nov 30;19(23):16047. doi:10.3390/ijerph192316047

6. Bruno F, Castelli G, Li B, Reale S, Carra E, Vitale F, et al. Genomic and epidemiological evidence for the emergence of a *L. infantum/L. donovani* hybrid with unusual epidemiology in northern Italy. Sibley LD, Sacks DL, editors. mBio. 2024 Jul 17;15(7):e00995–24. doi:10.1128/mbio.00995-24

7. Catta-Preta CMC, Sacks DL. Genetic Exchange in Leishmania: Understanding the Cryptic Sexual Cycle. Annu Rev Microbiol. 2025 Oct 23;79(1):105–28. doi:10.1146/annurev-micro-050724-091231

8. Rogers MB, Downing T, Smith BA, Imamura H, Sanders M, Svobodova M, et al. Genomic Confirmation of Hybridisation and Recent Inbreeding in a Vector-Isolated Leishmania Population. Didelot X, editor. PLoS Genet. 2014 Jan 16;10(1):e1004092. doi:10.1371/journal.pgen.1004092

9. Svobodová M, Alten B, Zídková L, Dvořák V, Hlavačková J, Myšková J, et al. Cutaneous leishmaniasis caused by Leishmania infantum transmitted by Phlebotomus tobbi. Int J Parasitol. 2009 Jan;39(2):251–6. doi:10.1016/j.ijpara.2008.06.016

10. Taddei R, Bregoli A, Galletti G, Carra E, Fiorentini L, Fontana MC, et al. Wildlife Hosts of Leishmania infantum in a Re-Emerging Focus of Human Leishmaniasis, in Emilia-Romagna, Northeast Italy. Pathogens. 2022 Nov 7;11(11):1308. doi:10.3390/pathogens11111308

11. Aoun K, Bouratbine A. Cutaneous Leishmaniasis in North Africa: a review. Parasite. 2014;21:14. doi:10.1051/parasite/2014014

12. Benikhlef R, Chaouch M, Abid MB, Aoun K, Harrat Z, Bouratbine A, et al. ITS1 and cpb genetic polymorphisms in Algerian and Tunisian Leishmania infantum isolates from humans and dogs. Zoonoses Public Health. 2023 May;70(3):201–12. doi:10.1111/zph.13016

13. Schönian G, Nasereddin A, Dinse N, Schweynoch C, Schallig HDFH, Presber W, et al. PCR diagnosis and characterization of Leishmania in local and imported clinical samples. Diagn Microbiol Infect Dis. 2003 Sep;47(1):349–58. doi:10.1016/S0732-8893(03)00093-2

14. González-de La Fuente S, Peiró-Pastor R, Rastrojo A, Moreno J, Carrasco-Ramiro F, Requena JM, et al. Resequencing of the Leishmania infantum (strain JPCM5) genome and de novo assembly into 36 contigs. Sci Rep. 2017 Dec 22;7(1):18050. doi:10.1038/s41598-017-18374-y

15. Späth GF, Bussotti G. GIP: an open-source computational pipeline for mapping genomic instability from protists to cancer cells. Nucleic Acids Res. 2022 Apr 8;50(6):e36–e36. doi:10.1093/nar/gkab1237

16. Kluyver Thomas, Ragan-Kelley Benjamin, Pérez Fernando, Granger Brian, Bussonnier Matthias, Frederic Jonathan, et al. Jupyter Notebooks - a publishing format for reproducible computational workflows. In: Positioning and Power in Academic Publishing: Players, Agents and Agendas [Internet]. IOS Press; 2016 [cited 2025 Nov 19]. Available from: https://www.medra.org/servlet/aliasResolver?alias=iospressISBN&isbn=978-1-61499-648-4&spage=87&doi=10.3233/978-1-61499-649-1-87 doi:10.3233/978-1-61499-649-1-87

17. McKinney W. Data Structures for Statistical Computing in Python. In. Austin, Texas; 2010 [cited 2025 Nov 19]. p. 56–61. Available from: https://doi.curvenote.com/10.25080/Majora-92bf1922-00a doi:10.25080/Majora-92bf1922-00a

18. Reback J, jbrockmendel, McKinney W, Bossche JV den, Augspurger T, Roeschke M, et al. pandas-dev/pandas: Pandas 1.4.2 [Internet]. Zenodo; 2022 [cited 2025 Nov 19]. Available from: https://zenodo.org/records/6408044 doi:10.5281/zenodo.6408044

19. Pedregosa F, Varoquaux G, Gramfort A, Michel V, Thirion B, Grisel O, et al. Scikit-learn: Machine Learning in Python. J Mach Learn Res. 2011;12(85):2825–30.

20. Waskom M. seaborn: statistical data visualization. J Open Source Softw. 2021 Apr 6;6(60):3021. doi:10.21105/joss.03021

21. Lex A, Gehlenborg N, Strobelt H, Vuillemot R, Pfister H. UpSet: Visualization of Intersecting Sets. IEEE Trans Vis Comput Graph. 2014 Dec 31;20(12):1983–92. doi:10.1109/TVCG.2014.2346248

22. Virtanen P, Gommers R, Oliphant TE, Haberland M, Reddy T, Cournapeau D, et al. SciPy 1.0: fundamental algorithms for scientific computing in Python. Nat Methods. 2020 Mar 2;17(3):261–72. doi:10.1038/s41592-019-0686-2

23. Wood DE, Salzberg SL. Kraken: ultrafast metagenomic sequence classification using exact alignments. Genome Biol. 2014 Mar 3;15(3):R46. doi:10.1186/gb-2014-15-3-r46 PubMed PMID: 24580807; PubMed Central PMCID: PMC4053813.

24. Wood DE, Lu J, Langmead B. Improved metagenomic analysis with Kraken 2. Genome Biol. 2019 Nov 28;20(1):257. doi:10.1186/s13059-019-1891-0 PubMed PMID: 31779668; PubMed Central PMCID: PMC6883579.

25. Sievert C. Interactive Web-Based Data Visualization with R, plotly, and shiny [Internet]. 1st ed. Chapman and Hall/CRC; 2020 [cited 2025 Nov 19]. Available from: https://www.taylorfrancis.com/books/9780429824210 doi:10.1201/9780429447273

26. Cokelaer T, Desvillechabrol D, Legendre R, Cardon M. “Sequana”: a Set of Snakemake NGS pipelines. J Open Source Softw. 2017 Aug 30;2(16):352. doi:10.21105/joss.00352

27. Dobin A, Davis CA, Schlesinger F, Drenkow J, Zaleski C, Jha S, et al. STAR: ultrafast universal RNA-seq aligner. Bioinformatics. 2013 Jan 1;29(1):15–21. doi:10.1093/bioinformatics/bts635

28. Liao Y, Smyth GK, Shi W. featureCounts: an efficient general purpose program for assigning sequence reads to genomic features. Bioinformatics. 2014 Apr 1;30(7):923–30. doi:10.1093/bioinformatics/btt656

29. R Core Team. R: The R Project for Statistical Computing [Internet]. 2021 [cited 2025 Nov 19]. Available from: https://www.r-project.org/

30. Love MI, Huber W, Anders S. Moderated estimation of fold change and dispersion for RNA-seq data with DESeq2. Genome Biol. 2014 Dec 5;15(12):550. doi:10.1186/s13059-014-0550-8

31. Benjamini Y, Hochberg Y. Controlling the False Discovery Rate: A Practical and Powerful Approach to Multiple Testing. J R Stat Soc Ser B Methodol. 1995;57(1):289–300. Located at: JSTOR

32. Aslett M, Aurrecoechea C, Berriman M, Brestelli J, Brunk BP, Carrington M, et al. TriTrypDB: a functional genomic resource for the Trypanosomatidae. Nucleic Acids Res. 2010 Jan;38(suppl_1):D457–62. doi:10.1093/nar/gkp851

33. Alvarez-Jarreta J, Amos B, Aurrecoechea C, Bah S, Barba M, Barreto A, et al. VEuPathDB: the eukaryotic pathogen, vector and host bioinformatics resource center in 2023. Nucleic Acids Res. 2024 Jan 5;52(D1):D808–16. doi:10.1093/nar/gkad1003

34. Bussotti G, Gouzelou E, Côrtes Boité M, Kherachi I, Harrat Z, Eddaikra N, et al. *Leishmania* Genome Dynamics during Environmental Adaptation Reveal Strain-Specific Differences in Gene Copy Number Variation, Karyotype Instability, and Telomeric Amplification. Tschudi C, editor. mBio. 2018 Dec 21;9(6):e01399–18. doi:10.1128/mBio.01399-18

35. Martin SH, Jiggins CD. Interpreting the genomic landscape of introgression. Curr Opin Genet Dev. 2017 Dec;47:69–74. doi:10.1016/j.gde.2017.08.007

36. Dumetz F, Imamura H, Sanders M, Seblova V, Myskova J, Pescher P, et al. Modulation of Aneuploidy in *Leishmania donovani* during Adaptation to Different *In Vitro* and *In Vivo* Environments and Its Impact on Gene Expression. Gull K, editor. mBio. 2017 Jul 5;8(3):e00599–17. doi:10.1128/mBio.00599-17

37. Prieto Barja P, Pescher P, Bussotti G, Dumetz F, Imamura H, Kedra D, et al. Haplotype selection as an adaptive mechanism in the protozoan pathogen Leishmania donovani. Nat Ecol Evol. 2017 Nov 6;1(12):1961–9. doi:10.1038/s41559-017-0361-x

38. Bussotti G, Piel L, Pescher P, Domagalska MA, Rajan KS, Cohen-Chalamish S, et al. Genome instability drives epistatic adaptation in the human pathogen *Leishmania*. Proc Natl Acad Sci. 2021 Dec 21;118(51):e2113744118. doi:10.1073/pnas.2113744118

39. Piel L, Rajan KS, Bussotti G, Varet H, Legendre R, Proux C, et al. Experimental evolution links post-transcriptional regulation to Leishmania fitness gain. PLoS Pathog. 2022 Mar;18(3):e1010375. doi:10.1371/journal.ppat.1010375 PubMed PMID: 35294501; PubMed Central PMCID: PMC8959184.

40. Bussotti G, Benkahla A, Jeddi F, Souiaï O, Aoun K, Späth GF, et al. Nuclear and mitochondrial genome sequencing of North-African Leishmania infantum isolates from cured and relapsed visceral leishmaniasis patients reveals variations correlating with geography and phenotype. Microb Genomics. 2020 Oct 1;6(10). doi:10.1099/mgen.0.000444

41. Antoniou M, Haralambous C, Mazeris A, Pratlong F, Dedet JP, Soteriadou K. Leishmania donovani leishmaniasis in Cyprus. Lancet Infect Dis. 2008 Jan;8(1):6–7. doi:10.1016/S1473-3099(07)70297-9

42. Ferreira TR, Inbar E, Shaik J, Jeffrey BM, Ghosh K, Dobson DE, et al. Self-Hybridization in Leishmania major. mBio. 2022 Dec 20;13(6):e0285822. doi:10.1128/mbio.02858-22 PubMed PMID: 36394334; PubMed Central PMCID: PMC9764971.

43. Akopyants NS, Kimblin N, Secundino N, Patrick R, Peters N, Lawyer P, et al. Demonstration of genetic exchange during cyclical development of Leishmania in the sand fly vector. Science. 2009 Apr 10;324(5924):265–8. doi:10.1126/science.1169464 PubMed PMID: 19359589; PubMed Central PMCID: PMC2729066.

44. Louradour I, Ferreira TR, Duge E, Karunaweera N, Paun A, Sacks D. Stress conditions promote Leishmania hybridization in vitro marked by expression of the ancestral gamete fusogen HAP2 as revealed by single-cell RNA-seq. eLife. 2022 Jan 7;11:e73488. doi:10.7554/eLife.73488 PubMed PMID: 34994687; PubMed Central PMCID: PMC8794473.

45. Louzada-Flores VN, Pescher P, Cokelaer T, Ferreira TR, Latrofa MS, Mendoza-Roldan JA, et al. Cross-subgenus hybridization between Leishmania and Sauroleishmania informs on parasite genomic compatibility and transcriptomic adaptation. BioRxiv Prepr Serv Biol. 2025 Mar 25;2025.03.25.645178. doi:10.1101/2025.03.25.645178 PubMed PMID: 40568159; PubMed Central PMCID: PMC12190823.

46. Gutiérrez-Corbo C, Domínguez-Asenjo B, Pérez-Pertejo Y, García-Estrada C, Bello FJ, Balaña-Fouce R, et al. Axenic interspecies and intraclonal hybrid formation in Leishmania: Successful crossings between visceral and cutaneous strains. PLoS Negl Trop Dis. 2022 Feb;16(2):e0010170. doi:10.1371/journal.pntd.0010170 PubMed PMID: 35139072; PubMed Central PMCID: PMC8827483.

47. Lypaczewski P, Thakur L, Jain A, Kumari S, Paulini K, Matlashewski G, et al. An intraspecies Leishmania donovani hybrid from the Indian subcontinent is associated with an atypical phenotype of cutaneous disease. iScience. 2022 Feb 18;25(2):103802. doi:10.1016/j.isci.2022.103802 PubMed PMID: 35198868; PubMed Central PMCID: PMC8841885.

48. Lypaczewski P, Matlashewski G. Leishmania donovani hybridisation and introgression in nature: a comparative genomic investigation. Lancet Microbe. 2021 Jun;2(6):e250–8. doi:10.1016/S2666-5247(21)00028-8

49. Romano A, Inbar E, Debrabant A, Charmoy M, Lawyer P, Ribeiro-Gomes F, et al. Cross-species genetic exchange between visceral and cutaneous strains of *Leishmania* in the sand fly vector. Proc Natl Acad Sci. 2014 Nov 25;111(47):16808–13. doi:10.1073/pnas.1415109111

50. Ferreira TR. At the genetic crossroads of Leishmania: Emerging hybrids reshaping disease patterns. PLoS Pathog. 2025 Jun;21(6):e1013213. doi:10.1371/journal.ppat.1013213 PubMed PMID: 40460155; PubMed Central PMCID: PMC12132935.

51. Späth GF, Piel L, Pescher P. Leishmania genomic adaptation: more than just a 36-body problem. Trends Parasitol. 2025 Jun;41(6):441–8. doi:10.1016/j.pt.2025.04.002

52. Kato H, Cáceres AG, Hashiguchi Y. First Evidence of a Hybrid of Leishmania (Viannia) braziliensis/L. (V.) peruviana DNA Detected from the Phlebotomine Sand Fly Lutzomyia tejadai in Peru. Pimenta PF, editor. PLoS Negl Trop Dis. 2016 Jan 6;10(1):e0004336. doi:10.1371/journal.pntd.0004336

53. Seblova V, Myskova J, Hlavacova J, Votypka J, Antoniou M, Volf P. Natural hybrid of Leishmania infantum/L. donovani: development in Phlebotomus tobbi, P. perniciosus and Lutzomyia longipalpis and comparison with non-hybrid strains differing in tissue tropism. Parasit Vectors. 2015 Nov 25;8:605. doi:10.1186/s13071-015-1217-3 PubMed PMID: 26608249; PubMed Central PMCID: PMC4660806.

54. Bussotti G, Li B, Pescher P, Vojtkova B, Louradour I, Pruzinova K, et al. *Leishmania* allelic selection during experimental sand fly infection correlates with mutational signatures of oxidative DNA damage. Proc Natl Acad Sci. 2023 Mar 7;120(10):e2220828120. doi:10.1073/pnas.2220828120

55. Iantorno SA, Durrant C, Khan A, Sanders MJ, Beverley SM, Warren WC, et al. Gene Expression in *Leishmania* Is Regulated Predominantly by Gene Dosage. Weiss LM, editor. mBio. 2017 Nov 8;8(5):e01393–17. doi:10.1128/mBio.01393-17

56. Leprohon P, Legare D, Raymond F, Madore E, Hardiman G, Corbeil J, et al. Gene expression modulation is associated with gene amplification, supernumerary chromosomes and chromosome loss in antimony-resistant Leishmania infantum. Nucleic Acids Res. 2009 Jan 9;37(5):1387–99. doi:10.1093/nar/gkn1069

57. Rogers MB, Hilley JD, Dickens NJ, Wilkes J, Bates PA, Depledge DP, et al. Chromosome and gene copy number variation allow major structural change between species and strains of *Leishmania*. Genome Res. 2011 Dec;21(12):2129–42. doi:10.1101/gr.122945.111

