## Supplementary Figures for "Genetic introgression and transcriptomic plasticity are associated with enhanced *Leishmania infantum* pathogenicity causing human cutaneous leishmaniasis in Tunisia"

[1]Institut Pasteur, Université Paris Cité, INSERM U1347, Unité de Parasitologie moléculaire et Signalisation, 25 Rue du Dr Roux, 75015 Paris, France; [2]Instituto Carlos Chagas, Fiocruz Paraná, Curitiba, Paraná, Brazil; [3]Institut Pasteur, Université Paris Cité, Bioinformatics and BiostatisticsHub, F-75015 Paris, France; [4]Centre National de Recherche en Génomique Humaine (CNRGH), Institut de Biologie François Jacob, CEA, Université Paris-Saclay, F-91057, Evry, France; [5]Laboratoire de recherche, LR 16IPT06 « Parasitoses médicales, Biotechnologies et Biomolécules », Institut Pasteur de Tunis, Université Tunis El-Manar, 13 Place Pasteur, Tunis, Tunisie.

**Keywords:** *Leishmania infantum*, Adaptation, Genomics, Transcriptomics, Hybrids, Post-transcriptional regulation

**Co-Correspondance:**

### SUPPLEMENTARY FIGURES:

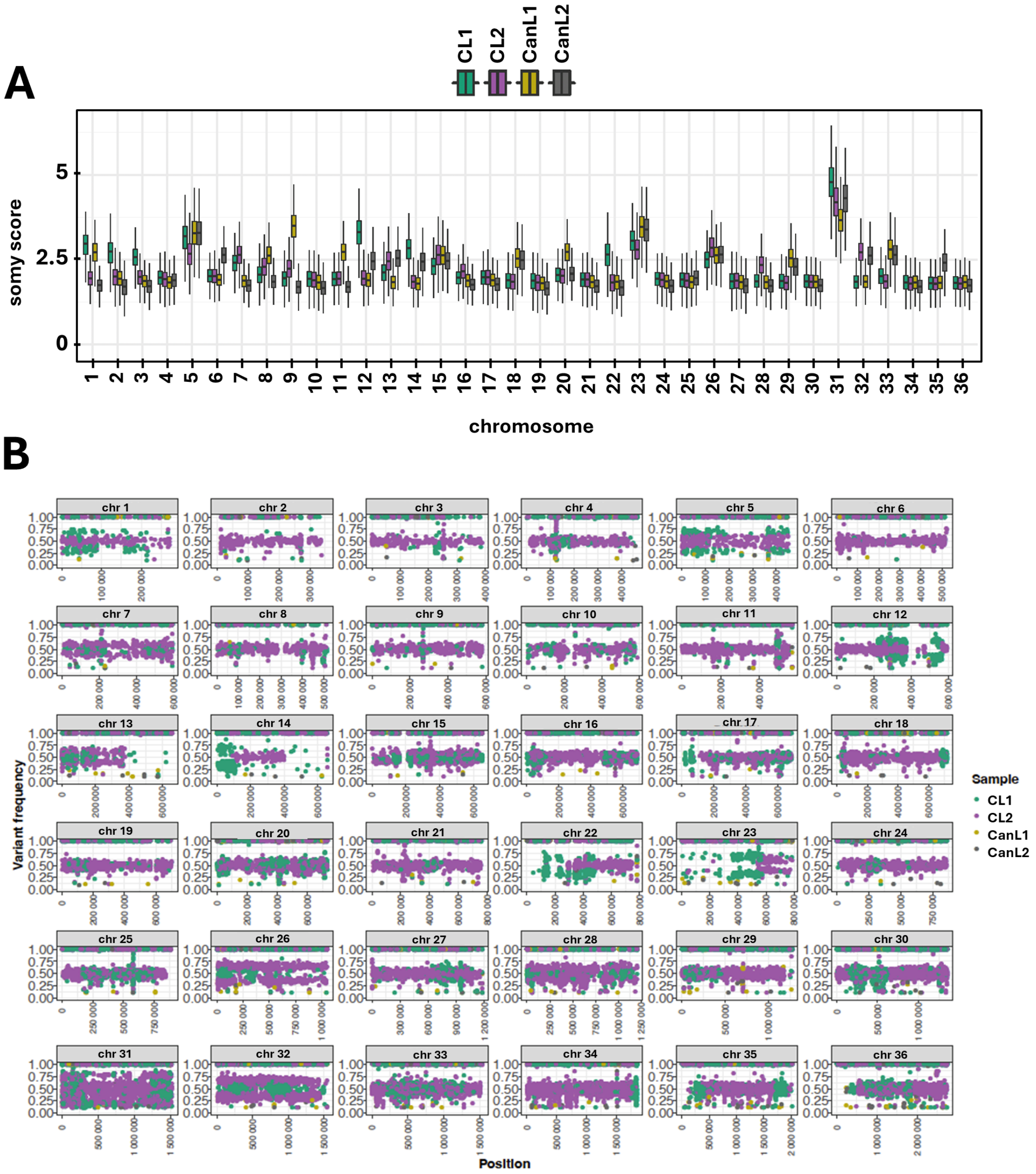


**Figure S1: (A)** Box plots indicating the copy number (expressed as somy score, y-axis) for the chromosomes (x-axis) across the 4 isolates indicated in the legend. **(B)** Dot plots indicating variant (SNP) frequeny (y-axis) as a function of genomic location (Position, x-axis) across the 4 isolates indicated in the legend.

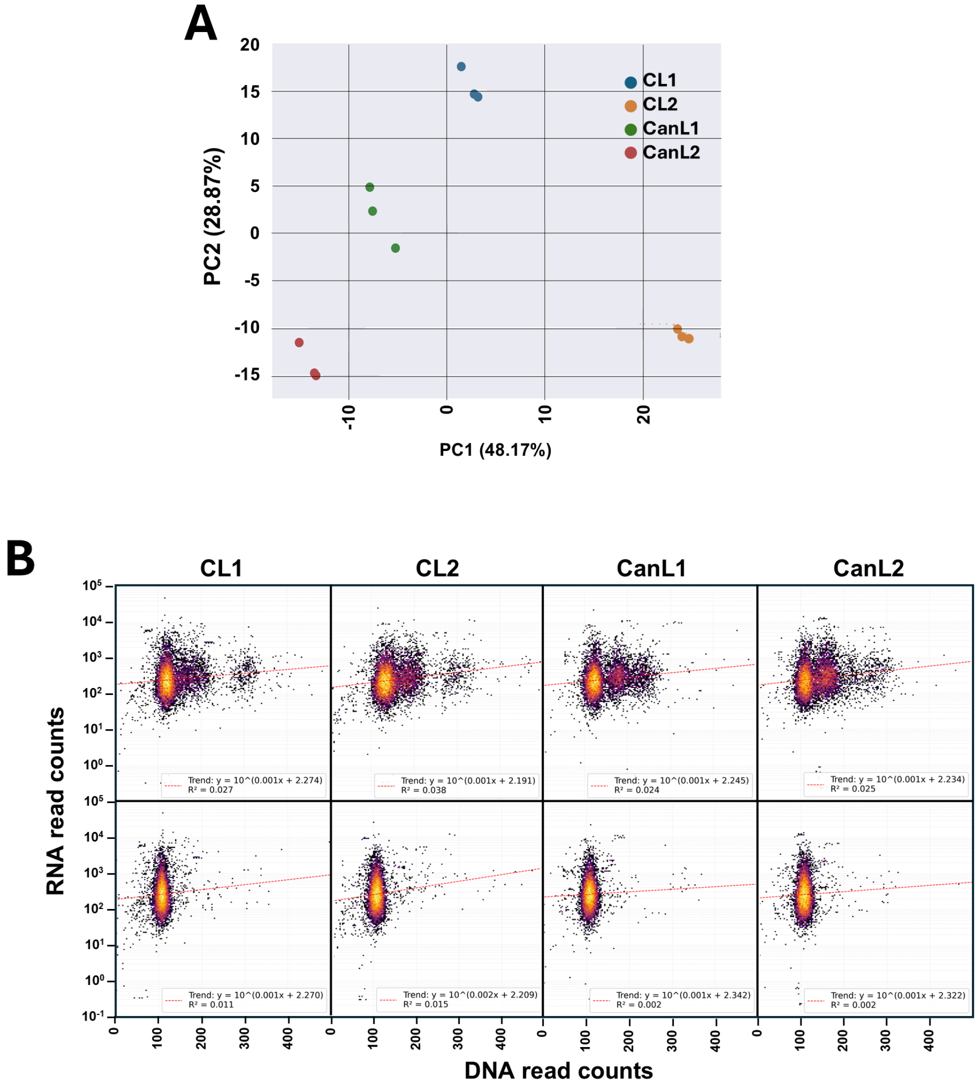


**Figure S2: (A)** Principal Component Analysis (PCA) of RNA-seq results obtained from three biological replicates of each sample. Samples are represented by coloured dots according to the shown legend. **(B)** Bin plots show the bivariate distribution and density of paired data points from the datasets RNA counts and DNA counts. Each bin aggregates data points falling within its area. The color of each bin represents the logarithm (base 10) of the number of data points it contains, revealing the densities. The red dashed line represents the linear regression trend line. RNA counts plotted against raw DNA counts (upper panel) or against DNA counts normalized for chromosomal somy (lower panel).
